# Availability of Medications Used for Puberty Induction and Maintenance in Adolescents with Hypogonadism in the Arab Region

**DOI:** 10.1101/2020.10.27.356808

**Authors:** Asma Deeb, Hussain AlSaffar, Rasha Tarif Hamza, Abdelhadi Habeb

## Abstract

**Purpose:** Inducing puberty in hypogonadal patients enables achieving normal final adult height, healthy bone mass accrual and improves fertility potential. Reliable availability and access to medicines remain a challenge around the world, particularly in low income countries. We aim to study the availability/access to medications used for inducing and maintaining puberty in centers within the Arab region.

**Patients and Methods:** A cross-sectional survey was conducted using a link to an online questionnaire which was emailed to paediatric endocrinologists in the Arab region. The questionnaire consisted of three questions related to availability of various forms of sex hormones.

**Results:** 99 physicians from 16 countries participated in the study. The commonest available form of estrogen was conjugated estrogen (29% of centers) followed by ethinylestradiol in 26%. Depot estradiol was available in 11centers while topical estrogen preparations of gel and patches were available in 6 and 10 centers respectively. Medroxy progesterone was available in 26% of the centers followed by noresthisterone (24%). The combined forms of oral and transdermal patches of estrogen/progestorne were available in 35 and 9% of centers. Intramuscular testosterone (Sustanon) was the most commonly available preparation of testosterone followed by the depot injection (Nebido), oral testosterone and testosterone gel and cream.

**Conclusions:** We report the first availability data of medications used for puberty induction and maintenance in paediatric hypogonadism in the Arab region. Recommended preparations for this purpose are not widely available. Creating essential list of medications used in paediatric endocrinology disorders might improve availability, access and consequently practice.

## Introduction

Paediatric endocrinology is a specialty with less common conditions and relatively higher cost medicines. Hypogonadism is a condition of sex hormone deficiency that results either from primary gonadal dysfunction or secondary hypogonadotropic causes.

In hypogonadal girls, estradiol deficiency causes endothelial dysfunction, reduced insulin production, abnormal lipid profile, increased central adiposity and formation of early atheroma (1). Inducing puberty enables patients to achieve normal final adult height and healthy bone mass accrual. It also alleviates psychological distress and improves fertility potential (2).

Estrogens, progestens and androgens are agents used to induce and maintain puberty. These agents have different pharmacokinetics and dynamics compared to endogenous steroids (3). World Health Organization (WHO) first created an Essential Medicines List (EML) for children in 2007, 30 years after the first EML for adults. Hormones are listed in the complimentary (rather than core) list of WHO essential medicines considering that they require specialized diagnostic, monitoring facilities, specialist medical care, and specialist training (4). To date, there are no clinical practice guidelines providing a master list of medicines used for paediatric endocrinology.

Reliable and sustainable availability and access to medicines remains a challenge around the world, with the poorest and most vulnerable populations at highest risk of failing to secure access. Up to a third of the population lack reliable sources for needed medicines. This proportion reaches higher than 50% in some countries in Asia and Africa (5). Some business model initiatives have been tried to improve access to essential medicines in limited resource countries and have shown positive impact on availability (6). However, these models were restricted to certain medications in limited areas.

Access to medicines that are used in high-income countries remains low in many low-middle income countries and prevents appropriate lifelong management of paediatric endocrine conditions (7). Medications listed in the complimentary essential list of WHO (like most of hormones) are usually linked to a higher price, which remains a major barrier for obtaining them.

Clinical practice guidelines in management of hypogonadism within paediatric endocrinology are variable. Choice of preparations used for inducing and maintaining puberty in males and females differ between Europe, the United States (US) and the Arab region (8, 9, 10). Lack of universal guidelines in management of hypogonadism might contribute to the limited availability of recommended preparations.

To the best to our knowledge, there is no data related to availability and access to medications used to induce and maintain puberty in hypogonadism within the Arab region.

## Aim

We aim to study the availability/access to medications used for inducing and maintaining puberty in female and male hypogonadism in centers within the Arab region.

## Materials and Methods

### Study design

This cross-sectional survey was conducted between July and October 2019 using a commercial software (Survey Monkey, USA). An invitation with a link to the online questionnaire was emailed to paediatric endocrinologists registered in the Arab Society of Paediatric Endocrinology and Diabetes (ASPED) database. This was followed by three reminders over the study period. The invitation outlined the purpose of the study, the voluntary nature of contribution, the unconditional right to decline participation and opting out from the database. Strict confidentiality of participants’ details was ensured and data were collected anonymously. The study was approved by the ASPED council. Considering that the study was survey-based utilizing a questionnaire for health care professionals without direct patient contact, research ethics approval was not deemed necessary. Health care professionals were asked to tick on a field of consent at the beginning of the survey confirming their willingness to participate in the study.

### The questionnaire

A comprehensive literature search on the study area was undertaken by the co-authors. A questionnaire was drafted in relation to availability of different forms of medications used for induction and maintenance of puberty in girls and boys. A hard copy of the draft was distributed during the 5^th^ ASPED-ESPE school in 2018 (11) and was validated by the school participants (12 consultants and 41 trainees). The final version of the questionnaire was approved by all co-authors and uploaded in the survey website.

The survey questionnaire was in English, which is the official language for educational communication between physicians in the region and included three questions. Different preparations of estrogen, progesterone and testosterone were listed in the three questions respectively. Participants were asked to select all available preparations in their centre of practice from the list and add ‘others’ as free text if not listed in the question **(Table 1)**.

**Table 1:** List of questions asked in the survey

| Which Preparation is Available in Your Centre? Please tick next to the ones available |  |  |
| --- | --- | --- |
| Question 1 | Question 2 | Question 3 |
| Testosterone | Estrogen | Progesterone & Combined Preparations |
| Intramuscular Testosterone (Sustanon) | Conjugated Estrogen | Norethisterone |
| Oral Testosterone Undecanoate | Ethinyl Estradiol | Utrogestan |
| Depot Testosterone Injection (Nebido/Reandron) | Estrified Estrogen | Medroxyprogesterone |
| Testosterone Cream 5% | Depot Estradiol Injection | Combined estrogen/progesterone pills |
| Transdermal testosterone gel | 17 $\beta$ estradiol | Combined estrogen/progesterone patches |
| Testosterone patch | Transdermal Estradiol Patch |  |
|  | Estradiol Gel |  |

## Results

Two hundred and seventy, three physicians opened the invitation email. Of those, 99 filled in the questionnaire giving a response rate of 36.3%. All 99 physicians answered the three questionnaire questions and indicated that they manage from five to over 20 patients with hypogonadism annually. The physicians were practicing in 99 different centers in 16 Arab countries; Morocco, Algeria, Tunisia, Libya, Egypt, Sudan, Jordan, Iraq, Lebanon, Palestine, Oman, Kuwait, Qatar, Kingdom of SaudiArabia (KSA) and the United Arab Emirates (UAE).

Forty nine centers were from the Gulf countries while 50 were from the North Africa and Levant countries. The highest response rate was from Oman (19%), followed by Saudi Arabia (16%) and Iraq (13%). Eight out of the 99 centers were private while the rest were governmental institutions (figure 1).

**Figure 1:**
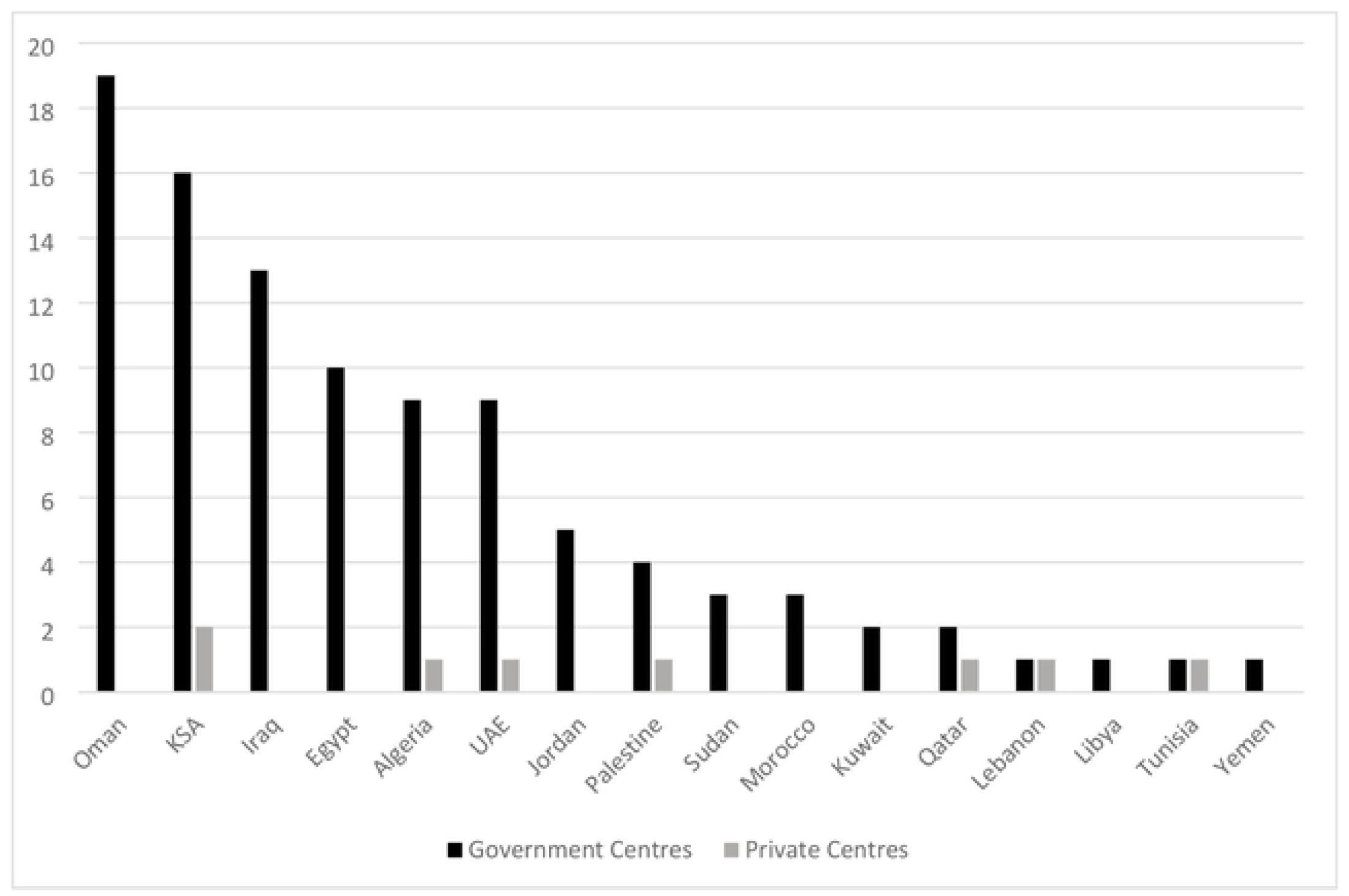
Centers of participating countries.

### Estrogen preparations availability

The commonest available form of estrogen was conjugated estrogen, which was available in 26 (29%) government and 3 (38%) private centers in 10 countries. This is followed by ethinylestradiol which was available in 24 (26%) and two government and private centers respectively from 11 countries. Estrified estrogen and 17 β estradiol were available in 15 and 11 centers in 8 and 6 countries respectively. Depot estradiol was available in 11centers in five countries. Topical preparations of estradiol, transdermal patches and estradiol gel were available in 10 and 6 centers in 8 and 4 centers respectively (Table 2).

**Table 2:** Available Estradiol Preparations within Countries and Centers. (G) indicates government centres

| Medicine | No. of Centers (G) | No. of countries | Which countries |
| --- | --- | --- | --- |
| Conjugated estrogen | 29 (26) | 10 | Algeria, Libya, Sudan, UAE, KSA, Iraq, Kuwait, Oman, Jordan, Qatar |
| Ethinylestradiol | 26 (24) | 11 | Algeria, Egypt, Sudan, Palestine, Jordan, Iraq, Qatar, Oman, Uae, KSA, Kuwait |
| Estrified Estrogen | 15 (13) | 8 | Tunisia, Algeria, Iraq, Jordan, KSA, Oman, UAE, Kuwait |
| 17 $\beta$ estradiol | 11 (8) | 6 | Algeria, Sudan, Tunisia, Iraq, UAE, Palestine |
| Depot Estradiol | 11(11) | 5 | Iraq, Qatar, Egypt, Jordan, Kuwait |
| **Transdermal**<br>**Estradiol patches** | 10 (9) | 8 | KSA, Oman, Iraq, Palestine, Egypt,<br>Algeria, Lebanon, KSA |
| **Estradiol gel** | 6 (4) | 4 | Algeria, Iraq, Oman, KSA |

### Progesteron and combined hormones preparations availability

Medroxy progesterone was available in 24 (26%) centers (2 private) in seven countries. This is followed by noresthisterone, which was available in 22 (24%) centers in eight countries while Utrogestan was available in eight centers in 3 countries. The combined forms of estrogen/progestorne enquired about were, combined contraceptive pills (OCP) and combined hormone patches. The former was available in 32 (35%) centers in nine countries while the latter was available in eight (9%) centers from 5 countries (Table 3).

**Table 3:** Available Progesterone and Combined Pills Preparation in Countries and Centers. (G) indicates government centres

| **Medicine** | **No. of**<br>**Centers (G)** | **No. of**<br>**countries** | **Which countries** |
| --- | --- | --- | --- |
| **Combined pills**<br>**(OCP)** | 32 (30) | 9 | Algeria, Egypt, Iraq, KSA, Oman, Kuwait, Qatar,<br>UAE, Sudan |
| **Oral:**<br>**Norethisterone** | 22 (21) | 8 | Egypt, Sudan, Libya, Iraq, Jordan, KSA, UAE,<br>Oman, |
| **Utrogestan** | 8 (6) | 3 | Algeria, Tunisia, UAE |
| **Medroxy P** | 24 (22) | 7 | Egypt, Sudan, Iraq, KSA, Oman, UAE, Kuwait |
| **Combined patches**<br>**(E+P)** | 8 (2) | 5 | Iraq, Palestine, Algeria, Oman, KSA |

### Testosterone preparations availability

Intramuscular testosterone (Sustanon) was the mostly available form of testosterone as it was available in all the 16 countries. Sixty one (67%) government and 8 (100%) private centers have access to this preparation of testosterone. Depot injection of testosterone (Nebido or Reandron) was available in 11(12%) government and one (12.5%) private centre in 6 countries (KSA, Egypt, Palestine, Yemen, Iraq and Oman). Oral testosterone preparation (testosterone undecanoate) was available in seven centers (one private) in five countries (KSA, Palestine, Oman, Iraq, UAE). Testosterone cream is available in five government centers in Iraq and Egypt while transdermal gel of 1 and 2% is available in 4 government centers in Iraq, Kuwait and Palestine. One government centre in Iraq had transdermal patch. Three participants from three government centers in Iraq and Egypt indicated availability of dihydrotestosterone (DHT) gel (Table 4).

**Table 4:** Available Testosterone Preparation in Countries and Centers. (G) indicates government centres

| **Medicine** | **No. of**<br>**Centers (G)** | **No. of**<br>**countries** | **Which countries** |
| --- | --- | --- | --- |
| **IM T (Sustanon)** | 69 (61) | 16 | All countries |
| **T depot**<br>**(Nebido/Reandron)** | 12 (11) | 6 | KSA, Egypt,<br>Palestine, Yemen,<br>Iraq, Oman |
| **Testosterone**<br>**undecanoate** | 7 (6) | 5 | KSA, Palestine,<br>Oman, Iraq, UAE |
| **Testosterone**<br>**cream 5%** | 5 (5) | 2 | Iraq, Egypt |
| **Transdermal gel: T**<br>**1%, 2%** | 4 (4) | 3 | Iraq, Kuwait,<br>Palestine |
| **T patch** | 1 (1) | 1 | Iraq |

### Overall preparations availability

The most commonly available preparation was intramuscular testosterone (sustanon) which was noted to be available in all countries and majority of the centers. Oral Estrogen preparations; conjugated estrogen and ethinyl estradiol were available in 29 and 26 centers of 10 and 11 countries respectively. Oral Medroxy progesterone and norethisterone were available in 8 countries (24 and 22 centers respectively). Testosterone undecanoate was available in 7 centers of 5 countries. In regards to the patches preparations, transdermal estrogen patches and combined estrogn/progesterone patches were available in 8 countries while testosterone patches were available only in one. Estradiol gel was available in 4 countries while testosterone gel of 1 and 2% were available in 3 countries (Figure 2).

**Figure 2:**
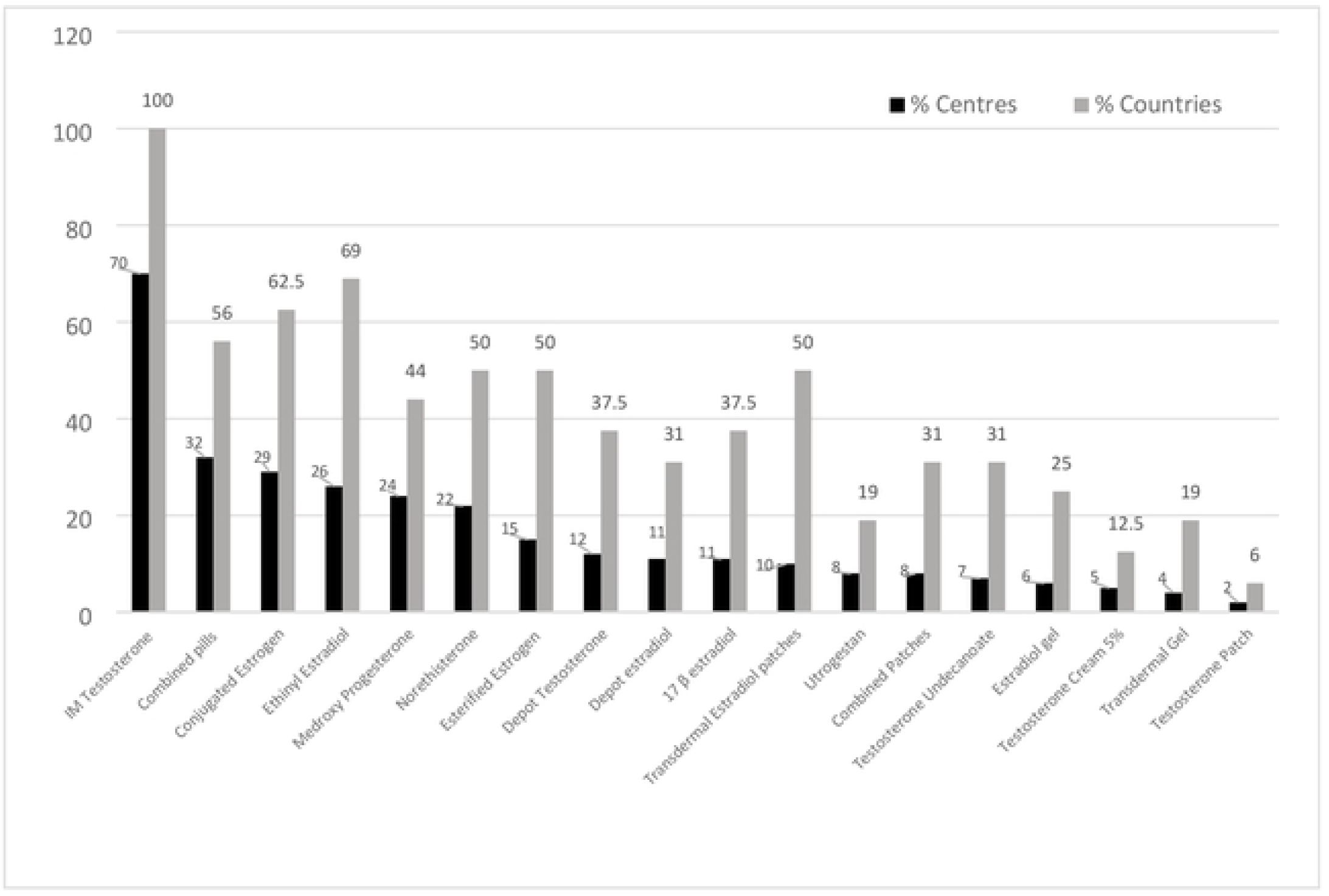
Overall list of medication in order of availability.

Comparison of individual preparation availability between Gulf and non − Gulf countries did not show any significant difference for any of the preparations.

## Discussion

In October 2003, WHO regional office in Cario conducted a pre-survey workshop on medicine availability and price measurement using the WHO/ Health Action International (HAI) methodology (12). It was followed by medicine prices survey conducted in nine Arab countries (Jordan, Kuwait, Lebanon, Morocco, Pakistan, Sudan, Syria, Tunisia and Yemen) (13). The workshop and the survey showed that availability of essential medications in public health facilities is generally suboptimal and needs improvement. Report showed that availability of medicines is lower in the public sector than in the private sector. Medicines available in the public sector are usually generics while those in the private sector is more likely to be of originator brands (12, 13). In this study, we were not able to study the difference of availability between government and private centers as the number of the latter was low.

Various explanations have been suggested for the variations in the selection of medication preparation to treat specific disease. This is applicable to the choice of hormone preparation used to treat hypogonadism. Availability and access are among factors influencing choice. In a previous study by our group, there was a difference in the selection of estrogen preparations by paediatric endocrinologists in Arab countries compared to USA and Europe (10). European and American guidelines recommend starting puberty induction with ethinylestradiol and conjugated estradiol respectively (8, 9). Both preparations were available only in around a quarter of the participating centre in this study. This might result in either not treating these patients with the resultant complication on bone and reproductive health or use of inappropriate available preparations. Estrogens are the first line of inducing puberty in girls with Progestogens introduced only after a period of unopposed oestrogen. Starting progestens without prior estorgenization results in uterine growth compromise particularly with preparations that have more androgenic effect such as norethisterone (14). Oral ethinyl estradiol was the most common form of estrogen prescribed by Arab physicians (29.7%) followed by oral conjugated estrogen (16.5%). Based on our current study, the varied selection cannot be attributed purely to availability as ethinylestradiol and conjugated estrogen were equally available in about third of participating centers (figure 2). The choice of one preparation or the other can be due to different prescription routines or training background in addition to availability. However, availability can be a deciding factor in choice of preparation. An example of the availability influencing choice is selection of oral 17 β-estradiol which was chosen by only 10% of practitioners in the Arab countries compared to 32% in Europe (8) and none in the US (9). The availability of oral 17 β-estradiol in participating centers is restricted to 11% which could be the reason for the low percentage of selection (figure 2). The use of estradiol patches is not high in our study or others. Patches were selected by 11.5% of Arab, 10% of European and 7.8% of US respondents (8-10) despite being more physiological. This could be explained by lack of availability of the preparation in our region. Less than 10% of participating centers have this form available (figure 2). There is increasing evidence that ethinylestradiol is inferior to transdermal oestradiol in terms of blood pressure and bone density which made transdermal oestradiol the preparation of choice (9). However, it was not a popular selection among Arab practitioners (10). This study confirmed that lack of availability of this preparation might be the main reason for not using it as patches were available only in one centre among the 8 private centers participated and only 9 out of the government centers.

80% of participants in our previous study (10) preferred IM testosterone esters. This choice was comparable to the US survey (9) where it was selected by 87.8% of responders. Androgen depot esters have been tried for over 50 years. They are known to be safe and have physiological pharmacokinetics. Intramuscular testosterone (Sustanon) was scored the highest available medication in our study (figure 2) with 69 (70%) centers from all countries stocking it. Long acting testosterone depot preparation is less commonly available with only 12% centers from six countries indicated its availability (table 1). In our previous study, there was a trend of using oral testosterone undecanoate in the Gulf states compared to other Arab countries (10). While we speculated the higher availability of this form in the Gulf might explain this trend, we could not confirm this speculation as we found no statistically significant difference in its availability between Gulf and non - Gulf countries (P = 0.2301).

Availability and selection of agents for treatment are inter-related issues. It is difficult to assume which results in the other. Ideally, a decision on choice is made by specialists on evidence-based grounds which, then ensures availability. This approach could be achieved by creating national essential medicines lists which can be country (or possibly region) specific. Lists need to address the disease burden of the nation/region, and include drugs used in various national health programmes. Creating these lists will not only reflect the relative disease burden but also highlight commonly used therapeutic interventions that needs to be made affordable, accessible, and available. These lists can be potentially powerful instrument to ensure that medicines are available, accessible and of good quality, resulting in strengthening health equity (15). Inclusion of more endocrine medications in WHO EMLs may promote the adoption of more medicines by individual countries within their national EML. To improve paediatric endocrinology medications availability, Rowlands et al recommended amending the cut-off age for children using physical maturity rather than the actual age. Also, to encourage formatting of the EMLs in a disease-focused manner rather than as individual medicines (16). On this issue, WHO ELM lists testosterone preparation as testosterone enanthate 200 mg/ml preparation, which is less frequently used compared to depot preparations. Listing the concentration of the preparation is another issue. For example, testosterone is listed as a 25 mg/ml testosterone propionate in the Indian national ELM when it is not the mostly used preparation nor the recommended dose/concentration (17). The same applies for estrogens where ethinylestradiol 0.01 and 0.05 mg is listed for use at primary, secondary and tertiary levels while lower dose estrogen pills suitable for younger girls with hypogonadism is not listed (17).

One of the main challenges in inducing puberty in girls is lack of licensed products with treatment frequently recommended by using off-label formulations intended for adults. In addition, while transdermal products are more physiological, they may be perceived as more difficult to use or less acceptable than oral products. Furthermore, there is very little information on specific dose response or bio-equivalency of oral or transdermal 17β oestradiol. These factors might influence prescription practice and consequently availability of these preparations. As for the progestens preparation, the recommended form is utrogestan 200mg once daily (18). This is a natural micronized progesterone which, provides good cycle control and is the least androgenic. Only eight centers out of 99 had this preparation available. Medroxyprogesterone was available in about a quarter of the centers from seven countries. Norethisterone was equally available but is not the preferred preparation to use considering its potent androgenic effect and its link to a higher incidence of dysmenorrhea (18).

In conclusion, we report the first data on the availability of medications used for puberty induction and maintenance in boys and girls with hypogonadism in the Arab region. Recommended preparations for this purpose are not widely available within the Arab region. Creating essential list of medications used in paediatric endocrinology disorders might improve availability, access and consequently practice.

## Disclosure

None of the authors has any conflict of interest to declare

## Funding

No funding was required

## Authors Contribution

All authors designed the study collectively. A Deeb coordinated the study tasks, supervised the overall work and wrote the initial draft. H Al Saffar managed the survey distribution and data analysis. RT Hamza and AM Habeb revised the manuscript.

